# PS membrane asymmetry influences the folding and insertion of a transmembrane helix

**DOI:** 10.1101/454504

**Authors:** Haden L. Scott, Frederick A. Heberle, John Katsaras, Francisco N. Barrera

## Abstract

The plasma membrane (PM) contains an asymmetric distribution of lipids between the inner and outer leaflets of its bilayer. A lipid of special interest in eukaryotic cells is the negatively charged phosphatidylserine (PS). In healthy cells, PS is actively sequestered to the inner leaflet of the PM but can redistribute to the outer leaflet when the cell is damaged or at the onset of apoptosis. The influence of PS asymmetry and its loss on membrane protein structure and organization have not been widely addressed. Marginally hydrophobic membrane proteins contain acidic residues in their transmembrane sequence, which can enable topological transitions after membrane association. The pH low insertion peptide (pHLIP), which undergoes a topological reorientation and inserts into the membrane at acidic pH – as its name implies, is a useful and well-characterized model for studying these transitions. Although it is known that the inclusion of PS in symmetric vesicles affects the membrane insertion process of pHLIP by lowering the pH midpoint of insertion, it is unclear how PS asymmetry influences these topological transitions. Here, we studied pHLIP’s topology using freely-floating asymmetric phosphatidylcholine (PC)/PS vesicles with PS enriched in the inner leaflet. We developed a modified protocol to create asymmetric vesicles containing PS and employed Annexin V labeled with an Alexa 568 fluorophore as a new probe to quantifying PS asymmetry. For pHLIP, membrane insertion was affected by the surface charge difference between bilayer leaflets caused by the asymmetric distribution of charged lipids between the leaflets. We thus conclude that lipid asymmetry can have consequences for the behavior of membrane-associated proteins. A corollary is that model studies using symmetric bilayers to mimic the PM may fail to capture important aspects of protein-membrane interactions.

## Introduction

The mammalian plasma membrane (PM) is a highly complex structure composed of hundreds of different lipids. Moreover, it has long been established that these lipids are not randomly arranged in the bilayer, but instead are asymmetrically distributed between the two bilayer leaflets (1). Specifically, the outer leaflet is enriched in phosphatidylcholines (PCs) and sphingomyelins, while the amino-containing glycerophospholipids such as phosphatidylserine (PS) and phosphatidylethanolamine (PE) are primarily located in the inner leaflet (1-4). Lipid asymmetry is entropically unfavorable and must be actively maintained by the cell. Two classes of ATP-dependent transporters, with the ability to move phospholipids unidirectionally to a given leaflet (5-8) have evolved for this purpose. These enzymes, termed flippases and floppases, counteract the movement of lipids down their concentration gradients by active translocation, however, the scrambling of the lipids occurs through the non-ATP dependent activation of the scramblase transporter (6, 9, 10). Although there is still considerable uncertainty regarding the physiological role of PM asymmetry, its importance for proper cellular function is not in doubt (9).

The maintenance of proper PS asymmetry is important for cellular viability (9). The presence of the negatively-charged PS in the inner leaflet enhances the binding of many cytosolic proteins, including key signaling proteins such as phospholipase C or KRAS (11, 12). Unsurprisingly, the loss of PS asymmetry is correlated with cellular malfunction and even death. For example, cell damage can result in the activation of scramblases, promiscuous lipid transporters that rapidly destroy membrane asymmetry (13). The resulting exposure of PS on the outer leaflet is a recognition signal for apoptosis to proceed via binding of annexins on the surface of macrophages (9).

Roughly half the mass of the membrane is proteins suggesting that membrane proteins are important constituents (14, 15). Transmembrane (TM) proteins contain sequences consisting largely of hydrophobic amino acids, driving protein insertion into the hydrophobic core of the membrane (16). Folding occurs through the formation of secondary structure that happens before membrane insertion (17, 18). It has been proposed that membrane proteins fold independently of their lipid environment (19). A recent study investigated the importance of the lipid environment using the bacterial protein LacY (20) using a recently developed method to create lipid asymmetry in freely floating model membranes (21). The authors found that PE asymmetry led to topological reorientations of LacY. However, the experimental system was far from ideal as LacY dissipated bilayer asymmetry by strongly increasing the rate of lipid flip-flop (20). That membrane asymmetry can influence the topological orientation of membrane proteins is worthy of further consideration. PM proteins are synthesized in the symmetric membrane of the endoplasmic reticulum (ER), yet their final destination is an asymmetric membrane (4, 22). It is unknown if the dramatically different ER and lipid environment from the ER to the PM affects membrane proteins that have the potential to change their topology after membrane insertion.

Here, we use the pH-low insertion peptide (pHLIP) as a model system to study how PM asymmetry affects the folding and insertion of membrane proteins. pHLIP assumes two different membrane topologies depending on the pH, as a result of changes in protonation in its seven charged residues. This characteristic enables experimental control of pHLIP’s topologies termed States II (membrane associated at neutral pH) and III (inserted as a TM helix at acidic pH) (23). Unlike membrane active peptides such as GALA, pHLIP does not lead to membrane leakage or disruption, a requisite for maintaining membrane asymmetry (23, 24). pHLIP has been studied in different membrane compositions and its behavior is well characterized (23, 25-27). Using symmetric PS vesicles, our lab previously showed that both the insertion p*K* and the insertion depth in State II decreased with increasing PS concentration (27). We proposed that both observations can be explained by an unfavorable interaction between the negatively charged PS head group and the seven negative charges present on pHLIP at neutral pH (27). However, it is unknown how a more biologically faithful model system, one in which PS is enriched in the inner leaflet, would influence pHLIP insertion. To test this, we modified a technique for producing freely-floating vesicles in order to mimic the asymmetric distribution of PS in the PM (28). We found that PS asymmetry caused an increase in the midpoint of pHLIP insertion, suggesting that for a physiologically relevant transbilayer charge distribution– i.e., less negative charge in the outer compared to the inner leaflet– lowers the energetic barrier for insertion. This finding proposes a general role for PS asymmetry in promoting the folding of TM proteins.

## Materials and Methods

### Materials

The lipids 1-palmitoyl-d31-2-oleoyl-*sn*-glycero-3-phosphocholine (POPCd31), 1-palmitoyl-2-oleoyl-*sn*-glycero-3-phosphocholine (POPC) and 1-palmitoyl-2-oleoyl-*sn*-glycero-3- phospho-L-serine (POPS) were purchased from Avanti Polar Lipids (Alabaster, AL) and used as is. Annexin V, Alexa Fluor 568 conjugate (Annexin V-568) was purchased form ThermoFisher Scientific (Waltham, MA) and assayed for concentration with an Agilent Cary 100 UV-Vis spectrophotometer (Santa Clara, CA) using an extinction coefficient of 23,380 M^-1^cm^-1^. pHLIP (sequence: N_t_-AAEQNPIYWARYADWLFTTPLLLLDLALLVDADEG-TCG-C_t_), synthesized using standard solid phase protocols and purified by reverse-phase high performance liquid chromatography (HPLC) to greater than 95% purity, was purchased from P3 Biosystems (Louisville, KY). A lyophilized pHLIP stock was dissolved in buffer (10 mM NaP_i_ pH 8.0) and assayed for concentration by UV-Vis using an extinction coefficient of 13,940 M^-1^ cm^-1^. Methyl-β-cyclodextrin (MβCD), sucrose, 4-(2-hydroxyethyl)-1-piperazineethanesulfonic acid) (HEPES), 2-(*N*-morpholino) ethanesulfonic acid (MES), calcium chloride (CaCl_2_), sodium acetate buffer (NaOAc), and sodium phosphate buffer (NaPi) were all purchased from Sigma-Aldrich (St. Louis, MO). Buffers were prepared by weighing the powder and adding ultrapure water to obtain the desired concentration. HEPES, MES, NaOAC were adjusted to the correct pH using hydrochloric acid or sodium hydroxide. Ultrapure water was obtained from a Millipore Milli-Q Academic (MilliPore Sigma, Burlington, MA) source.

### Asymmetric vesicle preparation and quantification

Asymmetric vesicles were prepared following the protocol from (28) with minor modifications due to the inclusion of POPS. Briefly, PS asymmetry was generated by using MβCD to catalyze the exchange of POPCd31 from multilamellar vesicles (MLVs) to the outer leaflet of large unilamellar vesicles (LUVs) composed of POPCd31 and POPS in a molar ratio of 93/7. Lipid films for donor MLVs were hydrated with 0.75 mL 20% w/w sucrose, and films for acceptor vesicles were hydrated with 0.5 mL ultrapure water; the concentrations of aqueous donor and acceptor were 25.3 mM and 12.6 mM, respectively. Donors were subjected to 4x freeze/thaw cycles while acceptors went through 5x cycles. Donor MLVs were diluted to 4.38 mM and incubated with 4.33 mL of 35 mM MβCD for two hours at room temperature in an 8:1 molar ratio. The acceptors were extruded through a 100 nm pore size polycarbonate membrane (31 passes) using a Mini Extruder (Avanti Polar Lipids, Inc., Alabaster, AL) to form large unilamellar vesicles (LUV). Acceptor LUVs were then added to the donor/MβCD mixture and incubated for one hour at 30°C, (although the original protocol calls for incubation at room temperature, we found that a slightly increased temperature allowed for greater PS exchange). The donor:acceptor molar ratio during exchange was 3:1. After the exchange step, residual MLVs were removed by centrifugation at 20,000 × g for one hour, and the supernatant containing the asymmetric vesicles was washed 4 times with buffer (1 mM HEPES/1 mM CaCl_2_/pH 7.4) and concentrated using a 100,000 MWCO centrifugal filter device (Amicon Ultra-15, EMD Millipore, Billerica MA). Gas chromatography coupled to mass spectrometry (GC/MS) (Agilent Technologies 7890A, Santa Clara, CA) was used to quantify the initial and final POPS concentrations in the vesicles after derivatization of the lipids to fatty acid methyl esters (FAMEs), as in (28). The FAMEs derived from protiated and deuterated palmitic acid were separately resolved by GC (29), allowing for quantitation of the mole fractions of POPCd31 and POPS (28). Three replicate samples of the initial acceptor vesicles and of the final asymmetric vesicles were measured to determine error bars. The mol % of POPS in the outer leaflet was calculated using the equation:

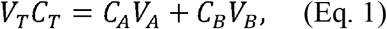

where *V_T_* is the total fractional vesicle volume (1), *C_T_* is the total concentration of POPS, *C_A_* and *C_B_* are the concentrations of POPS in the outer and inner leaflet, and *V_A_* and *V_B_* are the volume fractions of the outer and inner leaflet (0.51 and 0.49, respectively for 100 nm diameter vesicles). Vesicle concentration was determined by an inorganic phosphate assay (30). Dynamic light scattering (DLS) was used to determine the size and polydispersity of both the acceptors and asymmetric vesicles using either a Brookhaven Instruments BI-200SM (Brookhaven Instruments, Holtsville, NY) or a Wyatt DynaPro NanoStar (Wyatt Technology, Santa Barbara, CA) instrument. For the Wyatt instrument, the MW-R model was set for globular proteins and the Rg model was set for spheres. For the Brookhaven Instrument, 90-degree light scattering was measured using a 633 nm HeNe laser light source operated at 30 mW and a detector aperture of 200 μm. Data collection time was 4 minutes and the obtained autocorrelation curve was fit with both CONTIN and cumulant analyses. DLS measurements were taken at 25°C.

### Annexin V assay and lipid flip flop

Symmetric POPC vesicles with 0, 1, 2.5, 3.5, 5, 8, 10, 15, 20, and 50 mol % POPS were prepared in buffer (1 mM HEPES/1 mM CaCl_2_/pH 7.4) as described above. Annexin V-568 (stock concentration determined via UV/Vis) and buffer (25 mM NaOAc/1 mM CaCl_2_/pH 5.4) were added to the vesicles and incubated in the dark for 1 hour; the final lipid and Annexin V-568 concentrations were 50 μM and 0.52 μM, respectively, and the final sample pH was 5.5. Fluorescence measurements were made at room temperature with a Photon Technology International (Edison, NJ) Quanta Master fluorometer using the following instrument settings: excitation wavelength 579 nm, emission wavelength 601 nm, 15 s integration, 90° excitation polarization, 0° emission polarization, and 4.8 nm excitation and emission slit widths. Appropriate lipid blanks were subtracted in all cases, and changes in intensity were normalized to the POPC control. Symmetric samples of varying POPS concentration were used to determine a calibration curve that was fitted with the equation:

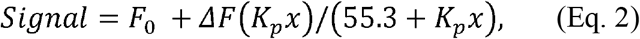

where *F_0_* is the initial fluorescence intensity, Δ*F* is the change in fluorescence intensity, *x* is the mol % of POPS, and 55.3 is the molar concentration of water (17, 31). This equation was used to determine the molar partition coefficient, K_p_, of Annexin V binding to the membrane in the presence of different levels of PS. Using this assay, we determined the molar % of POPS exposed at the outer leaflet of the asymmetric samples via a decrease in Annexin V-568 intensity as it bound PS. Using Annexin V-568 intensity values in the presence of asymmetric samples, we normalized the values to the PC control and inputted them into Eq. 2 to determine the mol % POPS in the outer leaflet. Measurements on asymmetric samples were taken for multiple days to assess the level of asymmetry and lipid flip flop.

Lipid flip flop in the presence of pHLIP was determined by performing the Annexin assay with asymmetric vesicles. pHLIP was incubated with POPC (control) and asymmetric vesicles for 1 hour at pH 7.4. Annexin V-568 was then added and incubated as described earlier. The final pH was 5.5, and the final pHLIP concentration was 0.25 μM. Intensity changes in the presence of pHLIP were analyzed as described earlier.

### Calculation of leaflet surface potentials

The surface potential of each leaflet in the asymmetric bilayer was calculated using the Grahame equation:

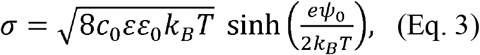

where σ is the surface charge density, *c*_0_ is the ion concentration, *ε* is the dielectric constant (here, 78.3), *ε*_0_ is the vacuum permittivity, *k_B_* is the Boltzmann constant, *T* is the absolute temperature (here, 298.2 K), and *ψ*_0_ is the surface potential (32). The surface charge density was calculated using the Guoy-Chapman Theory (32) separately for each leaflet using the measured POPS concentrations, and assuming an average area per lipid of 62.7 Å^2^ (33). Solving Eq. 3 for *ψ*_0_ gives the surface potential.

### Intrinsic tryptophan fluorescence spectroscopy

Symmetric POPCd31 vesicles with 0, 3, 7, and 30 mol % POPS were prepared *via* extrusion using a 100-nm pore size membrane (Whatman, UK) using a Mini Extruder (Avanti Polar Lipids, Inc., Alabaster, AL) in 10 mM NaP_i_ pH 8 to form LUVs. Asymmetric vesicles were in 1 mM HEPES/1 mM CaCl_2_/pH 7.4. Both symmetric and asymmetric vesicles were incubated with pHLIP for 1 hour at room temperature, for a final lipid:peptide molar ratio of 200:1. Final peptide concentration was 1 μM. It was previously shown that inclusion of the C-terminal cysteine in pHLIP does not cause disulfide-mediated dimerization (34). A pH titration was performed by adjusting the pH of the different samples with 100 mM stocks of NaOAc, MES, or HEPES buffers (25 μL), to obtain the desired final pH values. The final sample volume was 140 μL for symmetric samples, and 125 μL for asymmetric samples. A 2.5 mm-bulb pH-electrode (Microelectrodes, Inc., Bedford, NH) was employed to measure the final pH of each sample. Emission spectra were recorded using a Photon Technology International (Edison, NJ) Quanta Master fluorometer at room temperature with the following settings: excitation wavelength 280 nm, emission wavelength range 310-400 nm, and excitation and emission slits 3 nm. Appropriate lipid blanks were subtracted in all cases. Data were analyzed by calculating the spectral center of mass (CM) with the following equation:

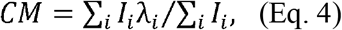

where *I_i_* is the fluorescence intensity at wavelength λ_*i*_. CM uses the entire spectral range of the data to inform on the local environment of the two Trp residues (35, 36). The data were also analyzed by monitoring changes in the fluorescence emission intensity FI at 335 nm, which is directly proportional to the population of molecular species present (37). CM and FI pH-titrations were then fitted to determine the p*K* using the equation:

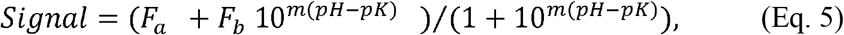

where *F_a_* is the acidic baseline, *F_b_* is the basic baseline, *m* is the slope of the transition, p*K* is the midpoint of the curve, and *Signal* refers to the changes in the fluorescence or circular dichroism signals as a function of pH.

### Circular Dichroism

CD measurements were performed using a Jasco (Easton, MD) J-815 spectropolarimeter at 25°C. pHLIP was incubated with POPC, symmetric POPC/POPS 97/3, or asymmetric vesicles (prepared as described earlier) in 10 mM NaP_i_ buffer (pH 8.0) for 1 hour. The pH was then adjusted with 100 mM NaOAc or NaP_i_ to a range of desired final pH values. The final sample volume, in each case, was 250 μL. For POPC samples, the lipid:peptide molar ratio was 200:1 with a final peptide concentration of 7 μM and a final lipid concentration of 1.4 mM. For the POPC/POPS 97/3 and asymmetric samples, the final peptide concentration was 3 μM and a final lipid concentration of 600 μM. Helical content changes of pHLIP were determined by measuring the ellipticity at 222 nm as a function of pH as in (38). Molar ellipticity was determined with the following equation:

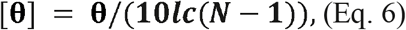

where Θ is the measured ellipticity, *l* is the cell path length, *c* is the protein concentration, and *N* is the number of amino acids (here, 38) (39). Calculated molar ellipticity at 222 nm was plotted against measured pH, and the resulting sigmoidal transition was fitted using Eq. 2 to obtain the p*K*_CD_. Spectra were collected from 260-195 nm for samples at pH 8 and 4 using the same temperature and scan rate as earlier, but with a 1 nm step size. Spectra were collected to check for secondary structure other than at 222 nm, which allowed for a detailed comparison of pHLIP in symmetric and asymmetric membranes.

### Statistical analysis

Statistical analysis was performed on the p*K* values and samples from the annexin assay to determine if the observed changes were significant. The analysis was performed using SPSSv25 software (IBM Analytics, Armonk, NY). One-way ANOVA and two different post hoc multiple comparisons tests were used based on the homoscedasticity (2-sided Dunnett t-test) or heteroscedasticity (Dunnett T3) of the data. Both the 2-sided Dunnett t-test and the Dunnett T3 set one variable as the control to compare to all other treatment groups. A 2-sided Dunnett t-test was used to determine the statistical significance of p*K* values determined via Trp fluorescence and CD by comparing all samples to the asymmetric vesicles sample. A one-way ANOVA was used to determine the statistical significance of the annexin assay results (Fig. S1). A Dunnett T3 test was used to determine the statistical significance of the stability of POPS asymmetry by comparing day 1 after the exchange to all subsequent days. A 2-sided Dunnett t-test was used to determine the statistical significance of lipid flip flop in the presence of pHLIP by comparing samples in the presence of pHLIP to samples without pHLIP. *P* ≤ 0.05 was considered significant for all tests.

## Results

### Preparation and quantification of asymmetric PS vesicles

The goal of this study was to understand how PS asymmetry influences the folding and insertion of a TM helix in a well-controlled and characterized model membrane system. A wide variety of techniques have been developed for the preparation of asymmetric bilayers, each with strengths and weaknesses (40, 41). For our study, it was important to use freely-floating vesicles, rather than solid supported bilayers, and to avoid the use of osmolytes that can potentially interact with the bilayer and/or create membrane tension (21, 42). To this end, we used the technique of methyl-β-cyclodextrin-mediated lipid exchange pioneered by the London group (21) with modifications that eliminate the requirement for concentrated sucrose in the vesicle core (Fig. 1) (28). Our strategy was to prepare symmetric POPC/POPS acceptor vesicles with PS as a minor component (~ 7 mol%), and then replace the POPS in the outer leaflet with POPC from a donor vesicle pool to generate asymmetric vesicles (aLUVs) with PS enriched in the inner leaflet.

**Figure 1.**
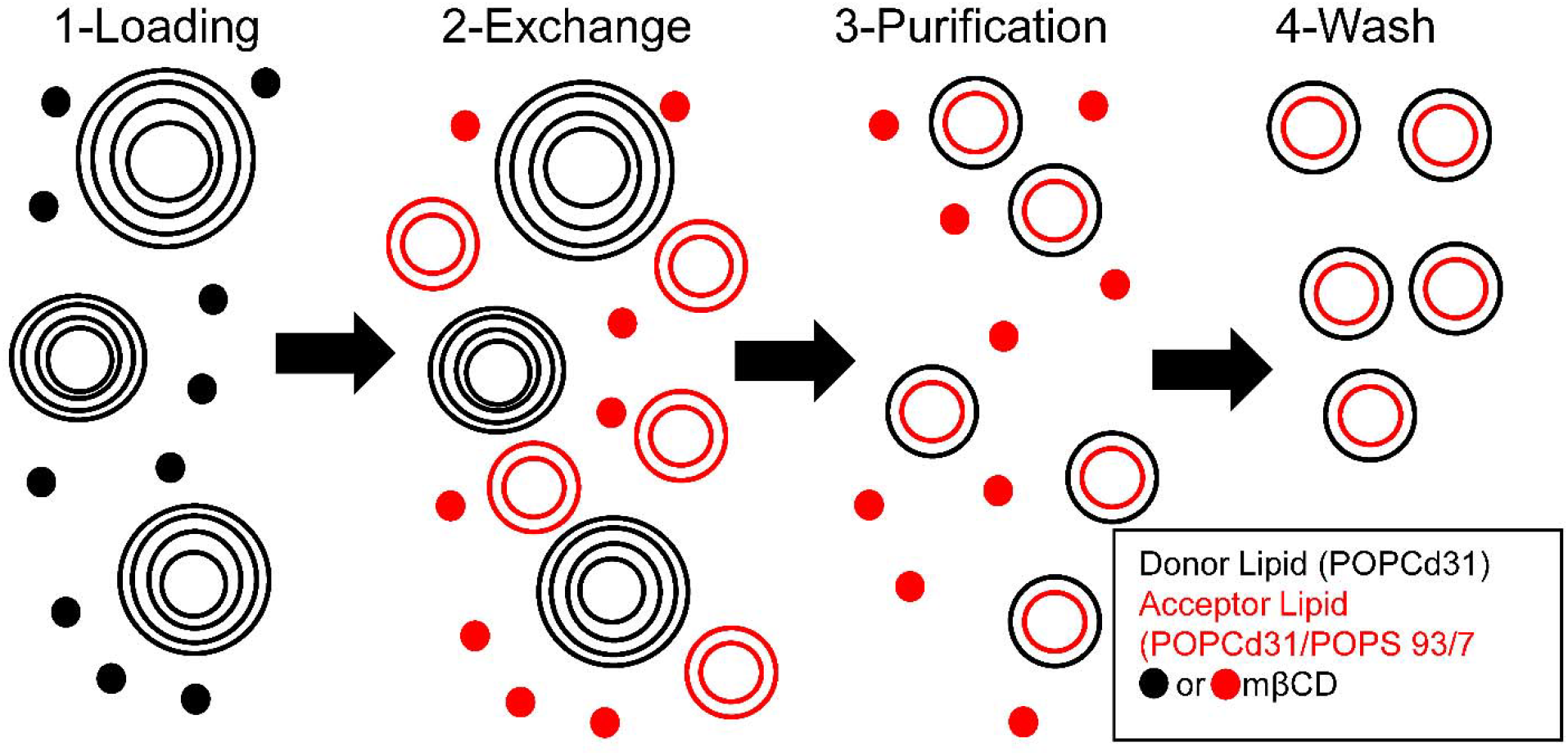
Cyclodextrin mediated exchange used to generate asymmetric vesicles. Steps were as follows: 1- mβCD is incubated with sucrose-loaded donor multilamellar vesicles to load donor lipid into mβCD. 2- Acceptor unilamellar vesicles are added for exchange to proceed with mβCD delivering donor lipid to the acceptors. 3- Donors are removed by centrifugation. 4- Asymmetric vesicles are washed in buffer of interest to remove mβCD. Black mβCD corresponds to the POPCd31-loaded state, and red mβCD is the loaded form containing POPS. Adapted from (28, 29).

aLUVs, once prepared, will gradually equilibrate to a symmetric state via passive lipid flip-flop. Although some studies have reported fast flip-flip (half times of seconds to minutes) in supported bilayers (43), there is general agreement that lipid flip-flop is much slower (half times of hours to days) in vesicles (6, 10, 44). To monitor PS asymmetry, we first used GC/MS to determine the total PS concentration in the aLUVs (28). Because GC/MS cannot distinguish between inner and outer leaflet lipid populations (28, 45), we employed externally added Annexin V conjugated with the Alexa-568 dye (Annexin V-568) to determine the amount of PS exposed in the outer leaflet. Annexin V specifically binds to PS headgroups in the presence of calcium (46) and is routinely used in cell biology for determination of the onset of apoptosis, which is marked by the exposure of PS to the extracellular environment (47). In symmetric vesicles, we observed a monotonic decrease in the fluorescence intensity of Annexin V-568 as PS concentration was increased (Fig. 2). Using this data as a calibration curve, we then assayed the concentration of exposed PS in aLUVs. Prior to outer leaflet exchange, the acceptor LUVs had an average exposed PS concentration of ~7 mol% (*N* = 5), which decreased to ~3 mol% in the aLUVs (Fig. S1). Statistical analysis revealed no significant differences in exposed PS concentration among the aLUV exchanges (*P* > 0.05), an indication that the preparation of asymmetric PS-containing vesicles was robust and reproducible.

**Figure 2.**
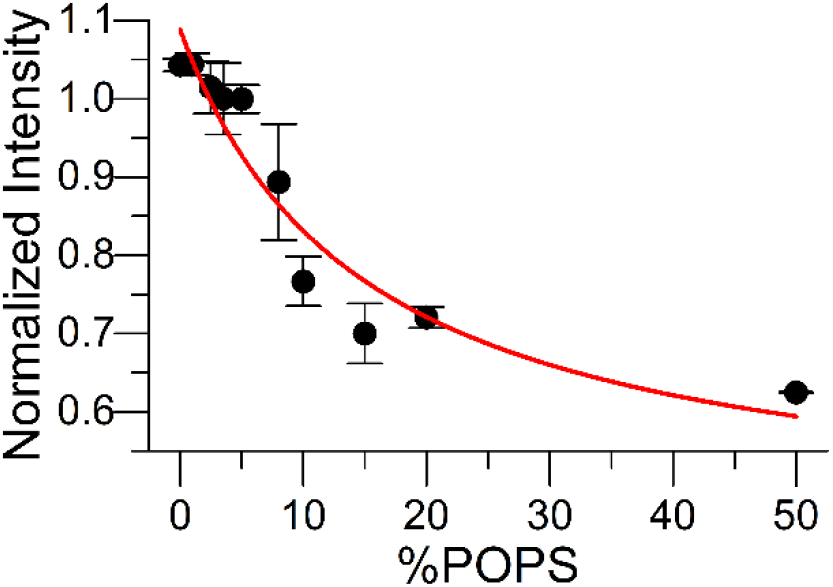
Annexin V-568 used to quantify the amount of PS in the outer bilayer leaflet. In the presence of PS, fluorescent Annexin V-568 exhibits decreased intensity with increasing concentration of exposed (outer leaflet) PS. Symmetric PC vesicles containing various levels of PS were used to generate a calibration curve. Red line is a fit to the data using a binding model (Eq. 2).

Next, we investigated the stability of aLUVs by monitoring PS asymmetry over time. Figure 3A shows that no significant loss of asymmetry in four-day old aLUVs occurred (*P* > 0.05). We also examined the influence of pHLIP on the stability of PS asymmetry. It has been previously reported that the presence of transmembrane peptides can accelerate lipid flip-flop and lead to a rapid loss of membrane asymmetry (48). However, we found that pHLIP in either State II or III did not lead to a significant loss of PS asymmetry over 3 hours via the annexin assay, a period of time greater than the time needed to complete our fluorescence spectroscopic assay (*P* > 0.05) (Fig. 3B). We also observed no changes in lipid flip-flop at room temperature over a period of 108 hours (data not shown). These results indicate that pHLIP does not influence the rate of POPS flip-flop on our experimental timescales.

**Figure 3.**
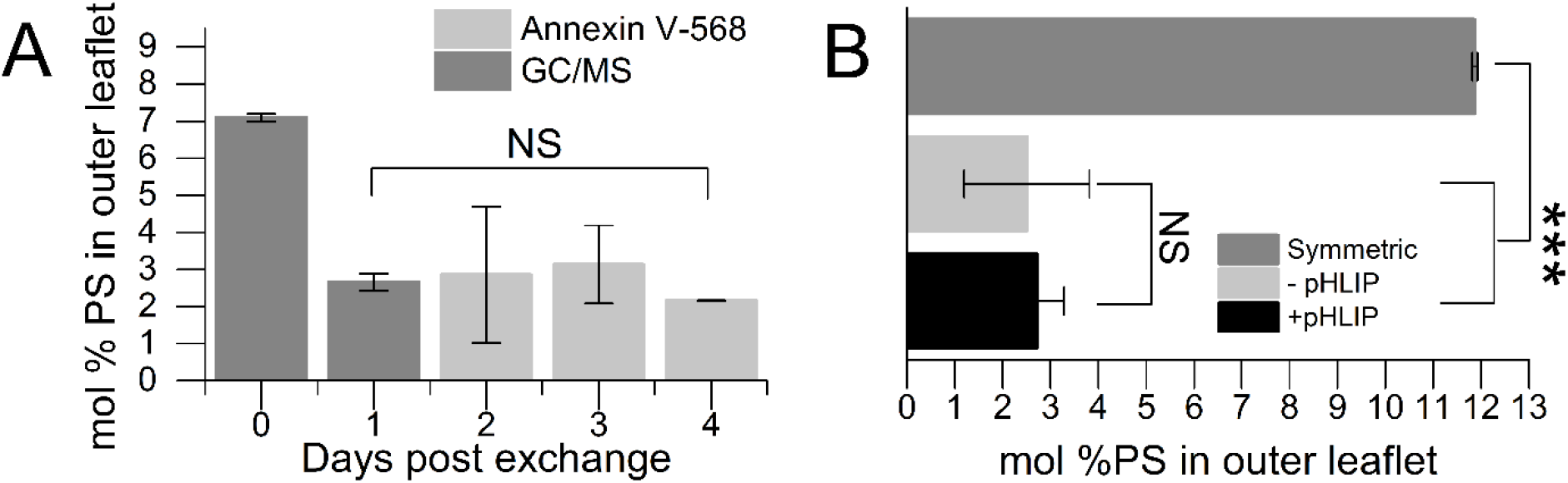
PS asymmetry is stable for days post exchange and in the presence of pHLIP. (A) Stability of asymmetric vesicles monitored for a period of four days post exchange. Day zero indicates the level of PS in the outer leaflet prior to exchange. Days 1-4 display the level of asymmetry post exchange. Both GC/MS and Annexin V-568 were used to determine the level of membrane asymmetry independently. Error bars represent the measurement uncertainty for each day obtained from replicates. (B) Comparison of PS levels in the outer leaflet of asymmetric vesicles incubated in the presence and absence of pHLIP. Here, pHLIP was incubated with asymmetric vesicles in both State II and III within the same sample by changing the pH after a given period of time. The symmetric vesicle PS level is prior to exchange and the asymmetric vesicle level is post exchange. Here the symmetric PS level was ~12%. The inclusion of pHLIP in the asymmetric vesicles does not lead to a loss of membrane asymmetry over a three-hour period post addition of pHLIP. ***, *P* ≤ 0.005; NS; no significance. Error bars are standard deviation.

### PS membrane asymmetry influences the insertion p*K* of pHLIP

The p*K* of insertion (midpoint of insertion) (23, 34) is a key parameter describing pHLIP’s interaction with membranes. pHLIP insertion occurs in multiple steps as pH is acidified (49, 50), and we recently reported that the insertion process is characterized by at least three macroscopic p*K* values that require different analyses and/or techniques for determination (38, 51). Specifically, when using the fluorescence emission spectra of pHLIP’s two Trp residues to monitor insertion into symmetric POPC vesicles, we found different insertion p*K* values depending on whether the response metric was the spectral center-of-mass (CM) or the fluorescence intensity at fixed wavelength (FI) (38). We refer to these insertion pH midpoints as p*K*_CM_ and p*K*_FI_, respectively.

We first used both native Trp and nitrobenzoxadiazole (NBD) conjugated to a C-terminal Cys residue fluorescence, as described previously (38), to determine pHLIP insertion p*K*s in symmetric POPCd31/POPS vesicles with increasing POPS concentration. As previously reported (52), decreasing the solution pH caused a shift in the Trp emission maximum to shorter wavelengths and an increase in intensity (Fig. 4A). These changes result from alterations in the local environment of the Trp residues, consistent with pHLIP insertion into the membrane. As NBD is also an environmentally sensitive dye, similar changes were observed compared to Trp (53-57). We then analyzed the emission spectra to determine the insertion p*K*_CM_, p*K*_FI_, and p*K*_NBD_ (Fig. 4B-E and Fig. S2). We found that p*K*_FI_ was ~0.5 units lower than p*K*_CM_ at each PS concentration and that both p*K*_CM_ (Fig. 4D) and p*K*_FI_ (Fig. 4E) decreased by ~ 0.4 units as POPS concentration increased from 0 to 7 mol %. Similarly, we observed a p*K*_NBD_ that was comparable to previous results, as well as a decrease in the presence of symmetric PS to a concentration of 7% (38). The comparable influence of PS suggests that deuteration does not influence the insertion of pHLIP. However, symmetric membranes do not recapitulate the PM, promoting the study of PS asymmetry.

**Figure 4.**
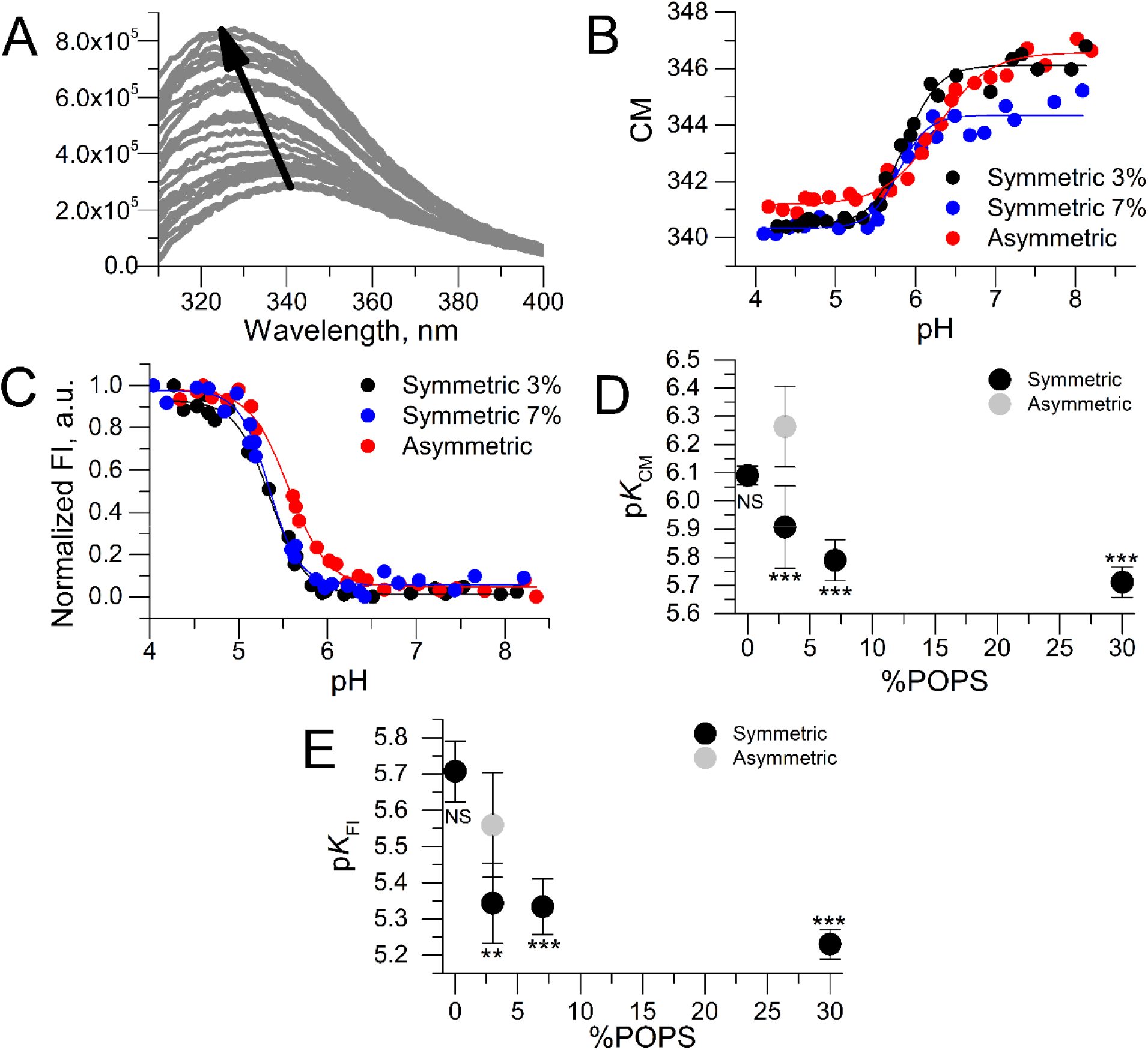
PS membrane asymmetry alters p*K_CM_ and* p*K_FI_*. Intrinsic Trp fluorescence indicates PS membrane asymmetry affects the p*K* of insertion of pHLIP causing a basic shift in the p*K* values compared to symmetric PS. (A) Representative Trp spectra displaying changes in Trp fluorescence as pH is decreased (arrow) in asymmetric vesicles. (B) and (C) Representative p*K* titrations for CM and FI Trp fluorescence analysis comparing asymmetric samples with symmetric samples with 3% and 7% PS. Titrations were fit to Eq. 3.to yield p*K_CM_* and p*K_FI_*. (D) and (E) shows a comparison of p*K* of insertion between asymmetric and symmetric vesicles with varying levels of PS for both a CM (D) and FI (E) analysis. Statistical analysis was conducted comparing asymmetric to symmetric samples (one-way ANOVA, 2-sided Dunnett t-test). **, *P* ≤ 0.01; ***, *P* ≤ 0.005; NS; no significance. Error bars are standard deviation.

We next examined aLUVs that typically contained ~3 mol % PS in the outer leaflet and ~7 mol % PS in the inner leaflet. We observed a statistically significant increase in both p*K*_CM_ and p*K*_FI_ compared to symmetric membranes with similar PS concentration (*P* < 0.05). Specifically, p*K*_CM_ of the asymmetric vesicles was 6.26 ± 0.14 compared to 5.91 ± 0.15 and 5.79 ± 0.07 for symmetric vesicles containing 3 and 7 mol % PS, respectively (Fig. 4D). Similar changes were observed in p*K*_FI_ – the aLUV value was 5.56 ± 0.14, while it was 5.34 ± 0.11 and 5.33 ± 0.08 for symmetric samples containing 3 and 7 mol % PS, respectively (Fig. 4E). Remarkably, p*K*_CM_ for the aLUVs was even higher than that of symmetric vesicles lacking PS, with the difference in the two p*K* values (i.e., p*K*_CM_ – p*K*_FI_) increasing to nearly 0.7 units, compared to 0.4-0.55 units for symmetric membranes. For p*K*_NBD_, we observed no statistically significant difference in the aLUV samples when compared to symmetric samples containing 3% PS. The increase in p*K*_CM_ and p*K*_FI_ suggests that PS asymmetry affects the membrane insertion of pHLIP in a manner that cannot be simply predicted from the outer leaflet composition.

### Secondary structure formation is not altered by PS membrane asymmetry

pHLIP must adopt a stable secondary structure prior to membrane insertion (17, 58). Using circular dichroism (CD), we previously observed that PS has no influence on the secondary structure of pHLIP in State II and III in symmetric vesicles composed of POPC/POPS 90/10 mol% (27). Figure 5A shows CD spectra of pHLIP in aLUV at pH values representing State II (pH 8) and State III (pH 4), revealing that pHLIP exhibits comparable helical content whether the membrane contains an asymmetric or symmetric PS distribution (27). This result suggests that PS asymmetry has little effect on pHLIP secondary structure at the initial and final states of the insertion process.

**Figure 5.**
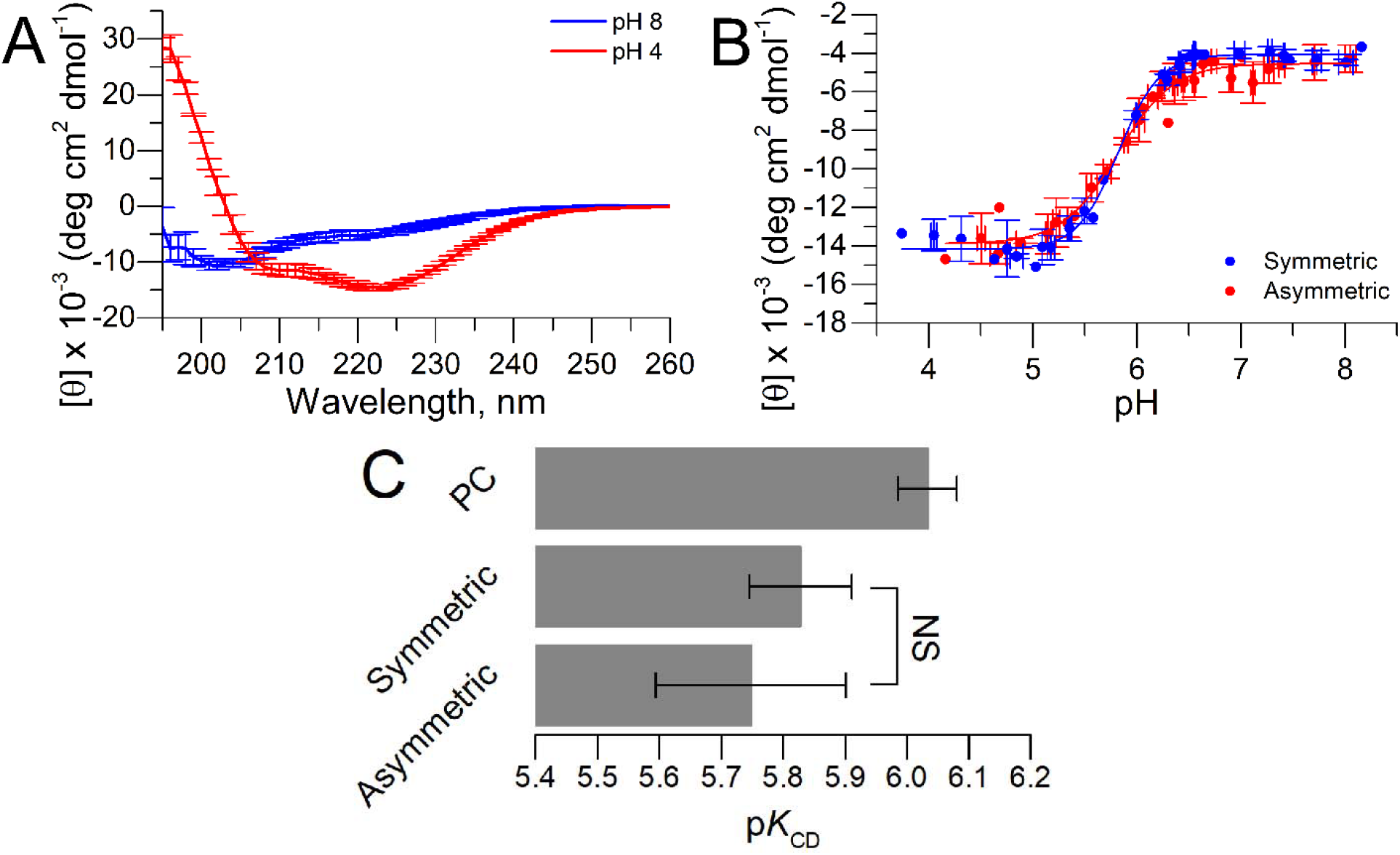
PS membrane asymmetry has no substantial influence on the helical formation of pHLIP. (A) Average CD spectra of pHLIP in the presence of PS asymmetry. pH 8 and 4 represent the membrane absorbed and transmembrane states of pHLIP, respectively. (B) Average CD titrations comparing symmetric and asymmetric POPC/POPS vesicles with 3% PS in the outer bilayer leaflets for both cases. Lines indicate fits to the data using equation 3. (C) p*K*_CD_ obtained from titrations of PC and symmetric PC/PS with 3% PS, and with asymmetric PS samples. No statistical significance (one-way ANOVA) is observed between symmetric and asymmetric PS, suggesting that asymmetric PS does not influence the insertion process of pHLIP as monitored by CD. NS; no significance.

As mentioned in the previous section, we recently proposed that different analysis methods for determining the insertion p*K* report independently on the protonation of different acidic residues in pHLIP (38). Specifically, we found that p*K*_CD_ informs on the midpoint of helical formation on the membrane surface. Since p*K*_CM_ and p*K*_FI_ showed a significant increase in asymmetric PS vesicles, we investigated if p*K*_CD_ was similarly affected. Figure 5B shows a comparison of average pH titrations of symmetric (3 mol% PS) LUVs and aLUVs containing ~ 3 mol% PS in the outer leaflet. Figure 5C shows no significant effect of PS membrane asymmetry on the p*K*_CD_ compared to symmetric samples (*P* > 0.05), suggesting that PS asymmetry affects only some steps in the membrane insertion of pHLIP.

## Discussion

### PS asymmetric vesicles can be prepared and are stable over multiple days

Membrane lipid asymmetry is a key property of cellular membranes (1). Membrane asymmetry is maintained by ATP-dependent enzymes that translocate lipids to their intended leaflet with high head group specificity, while subgroups of the ATP binding cassette proteins flop lipids with low head group specificity (5-8). Loss of these mechanisms of controlled lipid localization is a property of apoptosis and cell death (9). For example, Scott’s syndrome, a bleeding disorder, is associated with a problem in the regulation of membrane asymmetry, and is the only disease known to be with linked membrane asymmetry (59).

Here, we modified a recently developed technique that used cyclodextrin to create tensionless PC vesicles with an asymmetric chain distribution (28, 44). We aimed instead to create vesicles with an asymmetric PS distribution mimicking that found in the mammalian PM. We found that minor variations had to be made to the original technique to accommodate the movement of PS, possibly due to the slight differences in how mβCD solubilizes PS compared to PC (60). Once the vesicles were prepared, we needed to measure the level of PS asymmetry. A variety of techniques have been described for measuring the asymmetry of different lipid species and we first settled on nuclear magnetic resonance (NMR) as used by Heberle *et al.* for determining PS asymmetry (28). However, this approach failed, possibly due to a strong interaction between the positively charged chemical shift reagent Pr^3+^ and the negatively charged PS head group (data not shown). We also explored using trinitrobezenesolfonic acid (TNBS), a molecule known to interact with PE and PS (61). Unfortunately, TNBS showed little sensitivity to PS and failed to accurately detect PS levels in symmetric vesicles (data not shown).

After exhausting the available options from the literature, we took advantage of the well-known PS binding properties of Annexin V (46, 47), turning to a fluorescent version of this protein. We discovered that the fluorescence intensity of the Annexin V-568 conjugate is sensitive to the concentration of PS in the outer leaflet, which allowed us to assay for PS concentration in asymmetric vesicles using a calibration curve prepared from symmetric vesicles. We speculate the decrease in Alexa 568 fluorescent intensity is due to quenching caused by natural amino acids like Trp (62). Using this assay, together with GC/MS, we were able to consistently quantify the level of PS asymmetry in our aLUVs. The annexin assay showed that PS asymmetry could be generated and was stable for multiple days (Figs. 2 and 3). Using this same assay, we also determined that pHLIP did not disrupt asymmetry (Fig. 3B). This finding is of fundamental importance, since proteins and peptides often cause a loss of lipid bilayer asymmetry (20, 48), impeding studies looking into the effect of lipid asymmetry.

PS asymmetry may be particularly stable in part due to the unfavorable free energy barrier for transporting a negatively charged head group across the apolar membrane core. Moreover, pHLIP, with its many negative charges in State II, interacts adversely with PS (27). In State III, pHLIP does not disrupt the membrane and only interacts with a few lipid shells in close proximity to it (23, 63). This could explain our observation that pHLIP does not increase the rate of PS flip flop.

### PS asymmetry influences the insertion of pHLIP

The interaction of pHLIP with bilayers also depends on the specific lipid composition. However, all previous studies were carried out with symmetric bilayers (26, 27, 52, 64, 65). We have reported that symmetric PS vesicles decreased the insertion p*K* compared to PC vesicles, with a saturation at ~ 5 mol% PS (27). This was proposed to result from the unfavorable interaction between the negative charge on the PS headgroup and the seven negative charges on pHLIP at neutral pH (27). In a similar report, symmetric vesicles containing an assortment of lipid head groups, including PS, were also found to influence the insertion of pHLIP (64). However, an asymmetric distribution of PS, mimicking the plasma membrane, was missing from all these studies.

Here we study the effect of PS asymmetry on the membrane insertion of pHLIP. However, pHLIP’s membrane insertion is not fully described by one insertion p*K* and have been reported using different analysis methods (38). Specifically, in symmetric PC bilayers, p*K*_CM_ and p*K*_CD_ reported on the same pHLIP protonation event, whereas p*K*_FI_ reported on a different one (38). We found that the insertion p*K* in PS aLUVs determined from Trp fluorescence using both a CM and FI analysis is significantly increased (*P*<0.05) compared to a similar symmetric distribution of PS (Fig. 4). However, p*K*_CD_, which describes the midpoint of helical formation (38), was unaffected (*P*>0.05) (Fig. 5). Furthermore, we also determined that the p*K*_NBD_ (p*K* of translocation) was not influenced by PS membrane asymmetry compared to symmetric membranes with 3% PS. It should be noted that in our pH range used for determining the p*K*s, the PS headgroup remains deprotonated (66, 67). No changes in p*K*_CD_ or p*K*_NBD_ may indicate that PS asymmetry promotes the integration of Trp residues into the membrane before helical formation is complete, and that it does not influence the translocation of the C-terminus across the membrane (Fig. 6). As seen in Fig. 4 and 5, the titration reported by CM is largely complete before a large change occurs in the titration reported by CD, which indicates that in aLUVs CM and CD are decoupled in reporting the protonation of acidic residues (Fig. 6) (38). The decoupling of CM and CD suggests that PS asymmetry might change the p*K*a of D25 and D33 of pHLIP. Decoupling is not observed in samples containing symmetric PC or PS meaning that early protonations reported by CM are occurring before pHLIP begins to adopt its secondary structure in aLUVs. Equally important, the titration reported by FI only started at the point where the CM titration ends (Fig. 4) suggesting that CM and FI, as in symmetric POPC membranes, are reporting on different protonation steps in the insertion process (38). Together, the data indicates that PS asymmetry may influence the protonations of D14, 25, 31, and 33, as protonations of these residues are reported by CM and FI (38).

**Figure 6.**
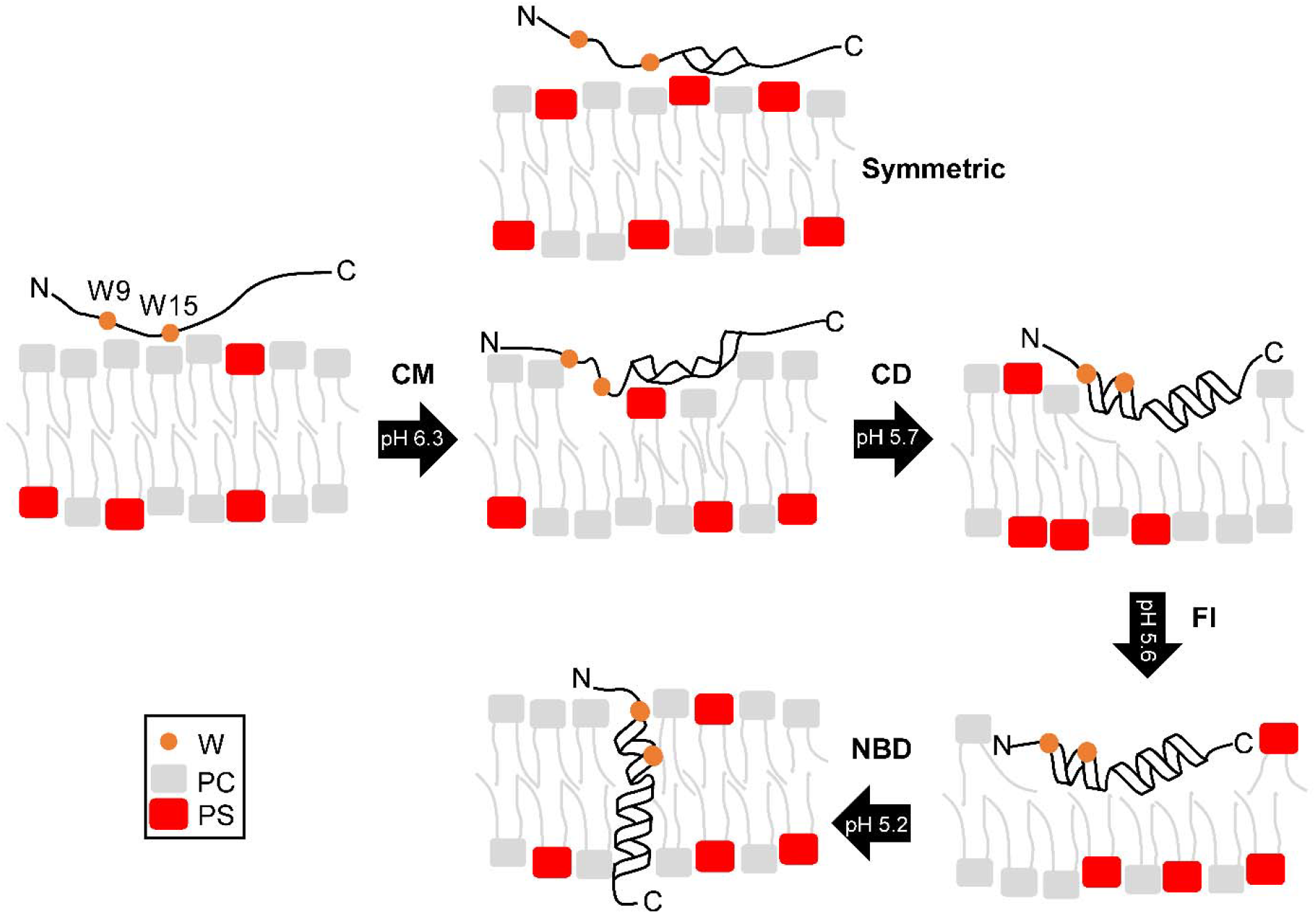
Influence of PS membrane asymmetry on the insertion of pHLIP. Black arrows are the transitions between each step in the insertion process, with the corresponding pH midpoints. Three transitions are represented by their insertion p*K.* The first transition represents the p*K* determined by CM. This transition is where the observed influences of membrane asymmetry are most prominent. Comparing to symmetric PS (directly above), the N-terminal region, including the two Trp residues, is more closely associated with the membrane in the presence of membrane asymmetry.

### Changes in membrane electrostatics promote an altered insertion of pHLIP

An asymmetric distribution of PS across the two leaflets is expected to change the electrostatics of the membrane (68-70). Lipid bilayers have three distinct potentials, namely the surface, transmembrane, and the dipole potentials (69). The surface potential is created by a buildup of charge on the membrane surface which propagates out from the membrane and into the aqueous solution (69). A concentration gradient of ions between the two aqueous solutions bathing the bilayer causes the transmembrane potential (69). Finally, the dipole potential is a positive potential centered in the hydrophobic core of the membrane, which arises from permanent dipoles in the lipid molecules (69). The observed increases in p*K*_CM_ and p*K*_FI_ could be caused by a change in one of these potentials (68-70). For example, an asymmetric distribution of negatively charged lipids should create a surface potential difference across the membrane (68, 70, 71). As a result, an additional transmembrane potential could potentially arise due to the asymmetric charge distribution from the PS head group. The membrane surface potential for each leaflet can in principle be calculated using the Guoy-Chapman model and the Grahame equation (Eq. 3) (32, 72). Although our protocol for creating PS asymmetry yielded only a small PS difference of ~ 4 mol%, this translates into an ~ +30 mV difference in surface potential across the membrane, with a less negative exterior. We propose that this surface potential might attract the negatively charged Asp and Glu side chains in pHLIP, leading to the peptide residing deeper in the interfacial region (68). Due to the higher dielectric constant and reduced hydration at this deeper location, the p*K*a values of these residues would be expected to increase (73). In contrast, p*K*_CD_ was not affected by PS asymmetry (Fig. 6), suggesting that asymmetry-induced changes in the surface potential do not affect the helical content of pHLIP (17) despite influencing its location at the membrane surface. Some caution with this interpretation is warranted, as surface potential differences may not be the only factor driving the observed p*K* shift. We cannot rule out that lipid asymmetry affects the lipid dipoles in a manner that alters the membrane dipole potential and thus influences the insertion p*K* of pHLIP (74, 75).

### pHLIP as a model can elucidate the influence of asymmetry on marginally hydrophobic TM domains

TM domains are primarily composed of hydrophobic amino acids, which anchor the domain into the bilayer. However, polar amino acids are commonly found in TM domains and indeed, Asp and Glu represent ~ 5% of the residues in TM sequences (16, 76). Typically, these polar residues are functionally important, such as in bacteriorhodopsin, where D85 and 96 (corresponding to D14 and 25 in pHLIP) mediate proton transfer across the membrane (77). A second example is the mammalian sodium/hydrogen exchanger NHE1, where conserved Asp residues located within the membrane environment allow this exchanger to control the internal pH of the cell (78). A more accurate amino acid sequence predictor of TM propensity discovered overlap between sequences that were both non-TM and TM (79). These sequences are termed marginally hydrophobic TM domains (mTMD) (80). mTMDs occur in membrane proteins and are unique in that they can assume two different topologies. These involve either movement of helices in and out of the bilayer, or reorientation of helices within the bilayer after insertion (81- 83). This orientation change is a direct consequence of the presence of polar residues, including Asp and Glu, which reduce their hydrophobicity compared to standard TMD (81-83). Some examples of proteins that have shown dual topology and mTMDs are the human aquaporin water channel APQ1, the hepadnaviral large envelope protein, and the ATP-gated ion channel subunit P2X2 (along with the related ASIC protein) (84-86).

Like mTMDs, pHLIP can adopt two different membrane topologies because of the content of its polar residues. In the case of pHLIP, the topological change can be easily measured in reconstituted systems, as TM insertion is triggered by a mere drop in pH. We propose that pHLIP might be used as a model system to gain mechanistic insights into topological transitions in membranes and speculate that membrane asymmetry might influence the membrane location of other peptides and proteins. Specifically, mTMD could experience topological transitions when exposed to membrane asymmetry changes, particularly those containing amino acids with charged side chains.

TM proteins are synthesized at the membrane of the endoplasmic reticulum and are transported to the PM *via* the Golgi apparatus. Membrane proteins experience environmental changes during this transit. Specifically, there is a difference in environmental pH, as the lumen of the Golgi is acidic, while the ER lumen and the cytoplasm have a neutral pH (87). However, it is poorly understood if such pH changes alter mTMD’s structure *via* changes to the protonation state of Asp, Glu or His residues. A second important consideration is that the ER and PM are different both in terms of lipid composition and asymmetry. Thus, the ER is delineated by a symmetric membrane, while the PM exhibits asymmetry of multiple lipid species, including PS (4). We have shown recently that the topology of pHLIP is affected by changes in symmetric lipid composition (27). Here, we show that not only the presence, but also their distribution in the bilayer impacts the location of pHLIP in the membrane. In the future work, we are interested in determining if asymmetry changes to the ER and PM can trigger mTMD topological changes, which could affect the folding and activity of membrane proteins in different cellular membranes.

## Author Contributions

H.L.S. performed the research. H.L.S., F.A.H., and F.N.B. designed the research and analyzed the data. H.L.S., F.A.H., J.K. and F.N.B. wrote the manuscript.

## Acknowledgments

We thank Dr. Drew Marquardt for thoughtful discussions about determining the level of PS asymmetry and his help with the NMR assays, Dr. Ilya Levental, Vanessa P. Nguyen, and Justin M. Westerfield for comments on the manuscript, and Forrest L. Davis for performing experiments. This work was partially supported by grant R01GM120642 from the NIH (F.N.B.), NSF grant No. MCB-1817929 (F.A.H.), and by funds from the UT-ORNL Joint Institute for Biological Sciences (JIBS) to F.N.B. Support was also received from the UTK-ORNL Science Alliance in the form of a Joint Directed Research and Development Award (to F.N.B.). aLUV preparation, GC/MS, and DLS measurements were supported by the Biophysical Characterization Laboratory suite of the Shull Wollan Center at Oak Ridge National Laboratory. J.K. is supported through the Scientific User Facilities Division of the Department of Energy (DOE) Office of Science, sponsored by the Basic Energy Science (BES) Program, DOE Office of Science, under Contract No. DEAC05-00OR22725.

